# A functional screening platform for engineering chimeric antigen receptors with reduced on-target, off-tumour activation

**DOI:** 10.1101/2020.04.30.070409

**Authors:** Raphaël B. Di Roberto, Rocío Castellanos Rueda, Samara Frey, David Egli, Rodrigo Vazquez-Lombardi, Sai T. Reddy

## Abstract

Chimeric antigen receptor (CAR) T cell therapies have advanced substantially in the clinic for cancer immunotherapy. However, challenges related to safety persist; one major concern is when CARs respond to antigen present on healthy cells (on-target, off-tumour response). A strategy to ameliorate this consists in engineering the affinity of CARs such that they are only activated by tumor cells expressing high antigen levels. Here, we developed a CAR T cell display platform for functional screening based on cell signaling. Starting with a CAR with high affinity towards its target antigen, we used CRISPR-Cas9 genome editing to generate a library of antigen-binding domain variants. Following multiple rounds of functional screening and deep sequencing-guided selection, CAR variants were identified that were discriminatively activated by tumor cells based on antigen expression levels. Our platform demonstrates how directed evolution based on functional screening can be used to enhance the selectivity and safety of CARs.

## INTRODUCTION

The clinical success of chimeric antigen receptor (CAR) T cells for cancer immunotherapy has demonstrated the potential of incorporating synthetic proteins in cellular therapeutics applications ^1^. CARs are hybrid proteins consisting of antigen-binding domains [e.g. antibody single chain variable chain fragments (scFv)] and intracellular signaling domains derived from the T cell receptor (TCR): the CD3 complex and receptors mediating T cell co-stimulation (e.g., CD3ζ, CD28, 4-1BB). Following viral delivery of CAR-encoding genes into T cells, the scFv enables recognition of tumour cells through surface antigen binding, while the intracellular signaling domains trigger the activation of a cytotoxic response. In a clinical setting, CAR T cells with specificity against the antigen CD19 have been successful in achieving partial and complete remission in patients with relapsed and refractory B cell leukemias and lymphomas ^2, 3, 4, 5^. As with other forms of adoptive T cell therapies, CAR therapies can be likened to “living drugs”, capable of achieving a sensitive, target-specific, self-amplifying and persistent response. But unlike cell therapies that rely on endogenous receptors, CARs benefit from their highly modular nature. For example, CAR specificity can be re-directed by engineering of the extracellular scFv domain without altering the other domains, thereby enabling targeting of a wide range of malignancies. Currently, clinical trials are underway to test the safety and efficacy of CARs against various tumour types and their antigens. These include cancers of the pancreas, liver, breast, gut and lung, all sharing tumour-associated antigens such as HER2, mesothelin, GD2 and CEA ^6^.

Although CAR therapies have shown clear clinical effectiveness against CD19-expressing B cell malignancies, other cancers have proven more challenging. Several pre-clinical and clinical trials have reported instances of on-target, off-tumour toxicities after CAR T administration ^7, 8, 9, 10^. Since healthy cells also have low-level expression of tumour-associated antigens (TAAs), CAR T cells can target such healthy tissue and lead to adverse events. This lack of specificity at the tissue level can be devastating for the organs affected and can require immediate treatment cessation due to the risk of patient fatality ^8^. In contrast, while the CD19 antigen is also present on healthy B cells, B cell aplasia resulting from CAR therapy is a manageable condition. The use of CAR T cells against non-CD19 cancers will thus require novel solutions in order to avoid serious adverse effects. HER2 is a prime immunotherapy antigen due to its overexpression in a wide range of cancers, most notably in mammary tumours ^11^, and its validity as a therapeutic target is well supported by the long success of the monoclonal antibody trastuzumab (Herceptin) in improving patient survival ^12^. While a T cell-based therapy targeting HER2 has the potential to induce a long term, persistent response, the downregulation of surface MHC-I in HER2-overexpressing cells makes this challenging ^13, 14^. CARs constitute an attractive alternative to the TCR, as they do not rely on MHC-based antigen presentation. Targeting HER2 with a CAR derived from the trastuzumab antibody was an effective strategy in a xenograft study ^15^, but proved fatal to a patient in a subsequent clinical trial ^8^. The respiratory distress and cytokine storm that followed CAR T cell injection were attributed to low levels of HER2 surface expression on normal lung epithelial cells.

Various strategies have been explored for improving the cell and tissue specificity of CAR T cell therapy. One common approach takes advantage of combinatorial antigen recognition, which involves the use of multiple receptors ^16, 17, 18, 19^ and/or soluble targeting molecules ^20, 21^ to induce logical decision-making in CAR T cells. Alternatively, the incorporation of suicide genes can be used to rapidly shut down a treatment gone awry ^22^. However, all these approaches require the genomic integration of multiple transgenes or a combination of transgenes and biologics, making an already complex therapy even more difficult to administer. Tuning the antigen binding affinity of a CAR scFv domain offers a simpler solution that is enabled by the existence of an apparent maximal T cell response above a receptor affinity threshold ^23^. This process typically involves selecting low affinity antibody or scFv variants from random or rational libraries and “exporting” the best candidates to a CAR format for further testing ^24, 25, 26, 27, 28, 29^. However, since the threshold for selectivity is unknown and may be context-dependent, many promising candidates fail once they are expressed as CARs.

Here, we report a T cell platform for CAR display that enables the engineering of variants based on antigen binding and/or signaling-based screening. Development of the platform required multiple steps of CRISPR-Cas9 genome editing, most notably the targeted integration of CAR genes into the unique genomic locus of the variable domain of the TCR β chain. As a measure of functionality, a green fluorescence reporter (GFP) reporter gene was integrated downstream of the endogenous interleukin-2 (IL-2) gene, facilitating high-throughput screening of activated T cells by fluorescence-activated cell sorting (FACS). We validated this functional screening approach by tuning the affinity of a CAR scFv domain with specificity towards the clinically relevant breast cancer antigen HER2. Starting with a scFv derived from the HER2-targeting therapeutic antibody trastuzumab, we generated a deep mutational scanning (DMS) library of CAR variants directly in our T cell platform via Cas9-mediated homology-directed repair (HDR). This library was then subjected to a series of iterative selection rounds based on IL-2 signaling activation following co-culture with a high-HER2-expressing cell line (SKBR3). For comparison, we also selected both weak and strong binders using soluble HER2 antigen. Deep sequencing was used to identify potential CAR variants that had lower binding affinity while maintaining similar signaling activation. Several variants from the signaling-based selection process showed enhanced discriminative recognition of target cells in a scenario mimicking on-target, off-tumour effects. These findings demonstrate the value of tuning CARs by using functional signaling-based screening and could be used as a general strategy for engineering CARs with both fine antigen specificity and target cell selectivity.

## RESULTS

### Genome engineering of a T cell platform for CAR display and functional screening

We generated a CAR display platform through a series of CRISPR-Cas9-based genome editing steps in the murine T hybridoma cell line B3Z ^30^. Previous work has shown that constitutive Cas9 expression in a cell line significantly enhances efficiency of non-homologous end joining (NHEJ) and HDR ^31^. Therefore, as our first step, we targeted the genomic safe-harbour locus *ROSA26* with exogenous Cas9 protein complexed with guide RNA (gRNA) and an HDR template encoding genes for Cas9 and blue fluorescent protein (BFP) under the control of the human CMV promoter (Figure 1a). This integration was confirmed by PCR amplification and Sanger sequencing (Supplementary Figure 1a, c). The functionality of endogenous Cas9 was subsequently affirmed through the NHEJ-mediated knock out of the BFP gene by transfecting BFP-targeting gRNA alone (Figure 1b).

**Figure 1:**
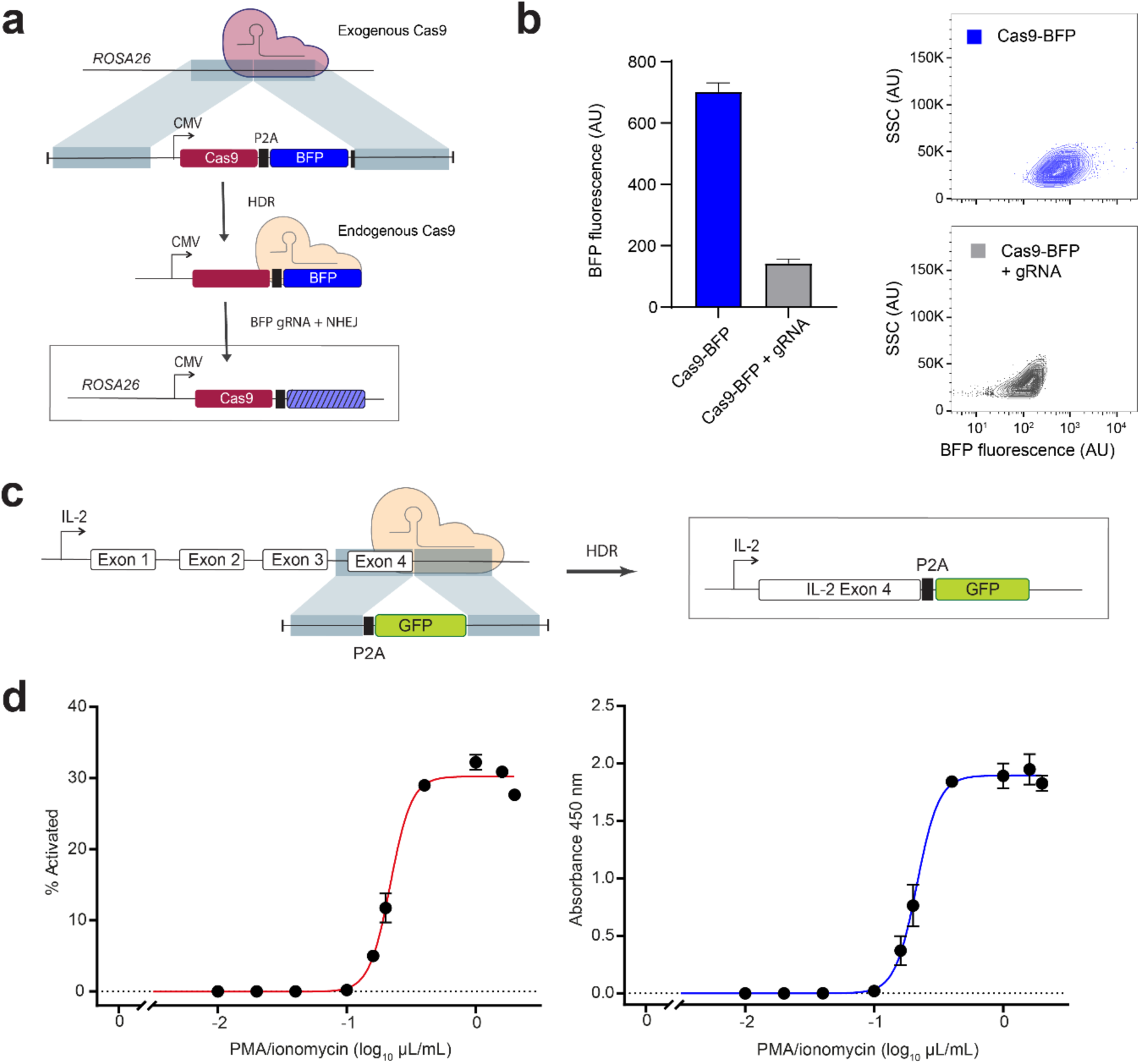
Engineering a reporter of IL-2 signaling in T cells by genome editing. **a** Integration of a Cas9 expression cassette in the genome of the B3Z T cell line at the safe harbour *ROSA26* locus using exogenous Cas9 protein and guide RNA (gRNA). The expression cassette consists of a gene encoding the Cas9 ORF, a P2A self-cleaving peptide sequence, and a BFP ORF under the control of the human cytomegalovirus (CMV) immediate-early enhancer and promoter. The constitutive expression of endogenous Cas9 was thereafter confirmed by the efficient disruption of the BFP ORF through the electroporation of BFP-targeting gRNA alone. **b** Histogram and flow cytometry plots showing difference in fluorescence between T cells expressing Cas9 before (blue population) and after (grey population) the knockout of BFP by gRNA electroporation. **c** Integration of a P2A peptide and GFP ORF immediately downstream of the final ORF of IL-2 in the genome of the Cas9-expressing B3Z cell line. **d** Dose-response curve of IL2-GFP expression in engineered T cells. Following overnight incubation with varying concentrations of the cell stimulation cocktail PMA/ionomycin (1X: 2 µL/mL), cells and culture supernatants were collected and assayed. The fraction of GFP-positive cells was measured by flow cytometry while IL-2 levels were measured by ELISA. The fraction of activated cells (red curve) matches the IL-2 secretion (blue curve) in terms of sensitivity, confirming that GFP fluorescence is a suitable reporter of IL-2 secretion.

IL-2 cytokine secretion is a reliable marker of T cell activation following antigen engagement and co-stimulation. Therefore, in order to generate a signaling reporter suitable for high-throughput screening, we used Cas9-mediated HDR to integrate a GFP open reading frame immediately downstream of the last exon of the endogenous IL-2 gene (Figure 1c), which was once again confirmed by PCR amplification and Sanger sequencing (Supplementary Figure 1b, d). An intervening P2A peptide ensures that GFP is not fused and secreted along with the cytokine. We confirmed by enzyme-linked immunoabsorbance assay (ELISA) that the genome edited T cells were still able to secrete IL-2 upon stimulation with a T cell activation cocktail of PMA/ionomycin. Interestingly, we found that the fraction of cells expressing GFP (not mean GFP intensity) scaled with cocktail concentration, and that the resulting dose-response curve matched well with that of IL-2 secretion (Figure 1d, Supplementary Figure 2). This binary response stands in contrast to previous work involving other T cell signaling reporters, such as nuclear factor of activated T cells (NFAT) ^32, 33, 34^, and may reflect our use of the endogenous IL-2 promoter. Going forward, we used the fraction of GFP-expressing cells as our T cell activation metric.

**Figure 2:**
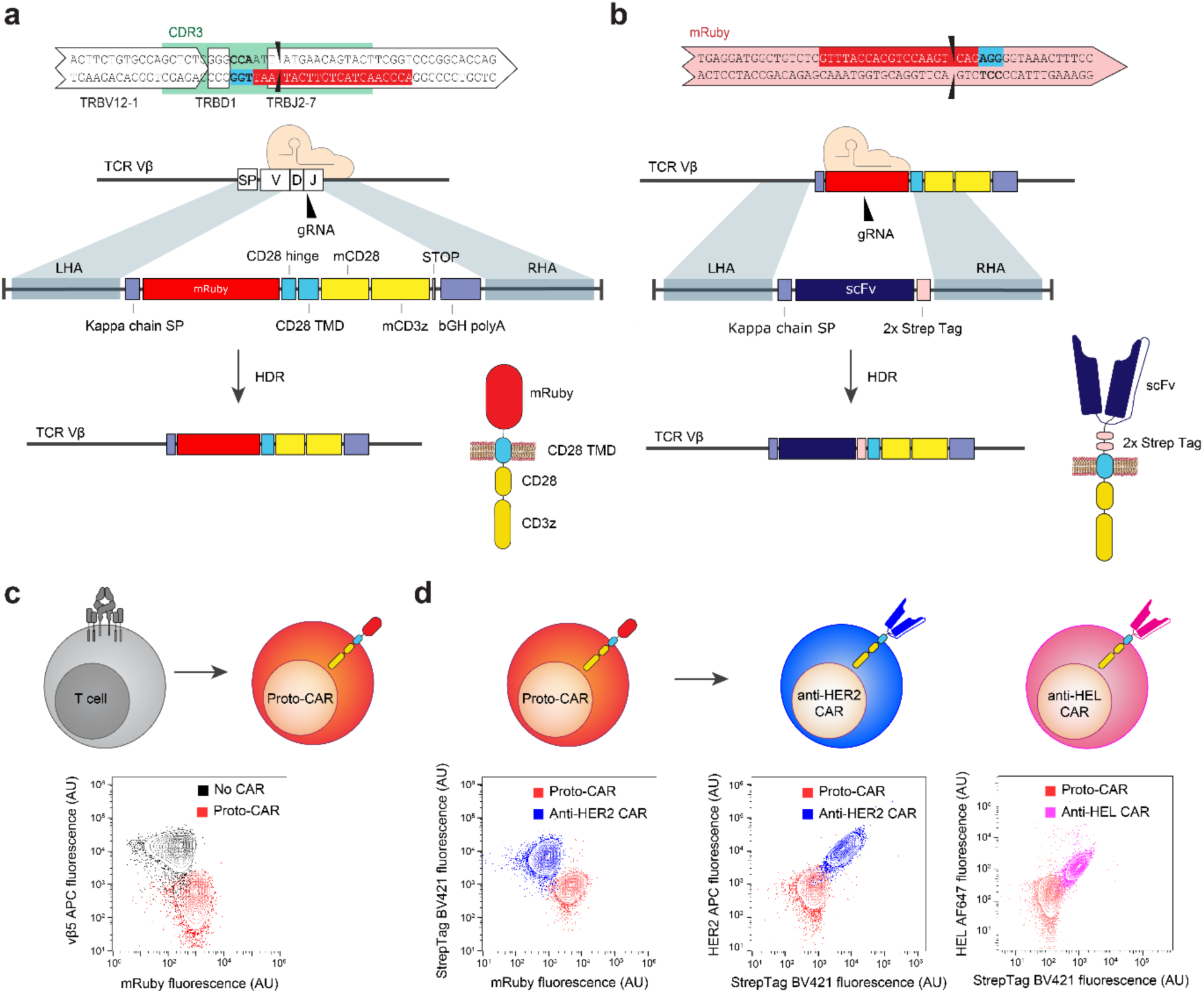
CAR expression and antigen binding in a T cell display platform. **a** CRISPR-Cas9 HDR is used for integration of a prototypical CAR (proto-CAR) in the unique CDRβ3 region of the TCRβ chain locus of engineered B3Z T cells. Double-stranded DNA with left and right homology arms (LHA and RHA) serves as repair template. A splice acceptor ensures that the CAR is on the same transcript and a signal peptide directs the protein to the secretory pathway. The proto-CAR includes a standard chassis with the CD28 hinge, transmembrane domain and co-signaling domain, and the CD3ζ signaling domain. The fluorescent protein mRuby is on the extracellular side of the CAR, as a placeholder for an scFv of choice. **b** CRISPR-Cas9 is used to replace the mRuby domain of the proto-CAR for a scFv and two Strep tags for detection. **c, d** Sequential genome engineering of T cell lines for the display of CARs with flow cytometry plots confirming CAR expression. The integration of the proto-CAR disrupts the surface expression of the endogenous TCR. This proto-CAR is then targeted to introduce an scFv of choice, such as one targeting the tumour antigen HER2 (blue cell schematic) or the model antigen HEL (pink cell schematic). The swapping of mRuby for an scFv can be detected by flow cytometry as the abrogation of mRuby fluorescence, the presence of a Strep tag II and the ability to bind labeled antigen.

Our next step was to genome engineer our reporter T cell line for CAR surface expression and signaling. The B3Z cell line expresses a TCR specific for an ovalbumin-derived peptide (SIINFEKL) presented on MHC-I ^30^. Therefore, we chose to target the TCR variable β chain complementarity determining region 3 (CDRβ3) for Cas9-mediated HDR of a CAR gene construct. Because of the diversity of V(D)J recombination, the CDRβ3 represents a unique genomic target site, which ensures that CAR expression is limited to a single variant per cell, eliminating confounding effects that can result from the multi-allelic or multi-site integration of different CAR variants. Targeting CAR integration into the Vβ chain locus also simultaneously ensures knock-out of the endogenous TCR. To provide a chassis for rapid CAR specificity changes, we first integrated a “proto-CAR” at the CDRβ3 locus. This CAR gene chassis is composed of the hinge and transmembrane domains of CD28 and the intracellular signaling domains of CD28 and CD3ζ, while lacking a typical binding domain, harbouring instead the fluorescent protein mRuby on its extracellular side (Figure 2a). The expression of mRuby provides a fluorescence reporter that can be used to easily detect HDR ^35^. We first confirmed integration by PCR amplification and Sanger sequencing (Supplementary Figure 1e) and then by flow cytometry we confirmed that this proto-CAR was expressed and that TCR expression was abrogated (Figure 2c). Next, we electroporated cells with gRNA targeting mRuby and HDR templates encoding scFv genes specific against the model antigen Hen Egg Lysozyme (HEL) (variant M3 ^36^) or human breast cancer antigen HER2 (variant 4D5/trastuzumab ^37^) (Figure 2b). Surface expression of CARs was confirmed by the presence of two Strep tags in the linker region between the scFv and the CD28 hinge domain. The ability of the CARs to bind their respective target antigens was also confirmed using soluble fluorescently-labeled cognate antigens (Figure 2d).

### CAR T cell signaling is activated with both high and low HER2-expressing cells

We next determined if our CAR T cell display platform expressing the anti-HER2 scFv could be activated following co-culture with HER2-expressing cell lines. We first selected a panel of cancer cell lines known to express different levels of HER2 on their surface ^38^. Flow cytometry confirmed that SKBR3, MCF-7 and HEK293 cells express decreasing levels of surface HER2 (Figure 3a). These cell lines therefore provide a model system for evaluating on-target, off-tumour toxicity of CAR T cells whereby SKBR3 represent tumour cells (high expression of HER2) and MCF-7 and HEK293 mimic off-target healthy cells (low expression of HER2). These cell lines were then co-cultured with CAR T cells expressing anti-HER scFV or anti-HEL scFv (negative control) and activation was measured through IL-2-linked GFP expression via flow cytometry (Figure 3b and c). Correlating with the amount of HER2 expression on the different cell lines, we observed substantial anti-HER2 CAR T cell activation in the presence of SKBR3 and MCF-7 cells, but not with HEK293, while anti-HEL CARs failed to elicit significant T cell activation with any co-cultured cells. We also tested a CAR based on the phage-derived scFv F5 for which affinity is estimated to be 275-fold lower than 4D5 ^26, 39^ but it was shown to be largely unresponsive to our cell lines. Interestingly, T cell activation appeared to correlate with trogocytosis, the transfer of antigen from target cells to lymphocytes, which is consistent with a recent study ^40^.

**Figure 3:**
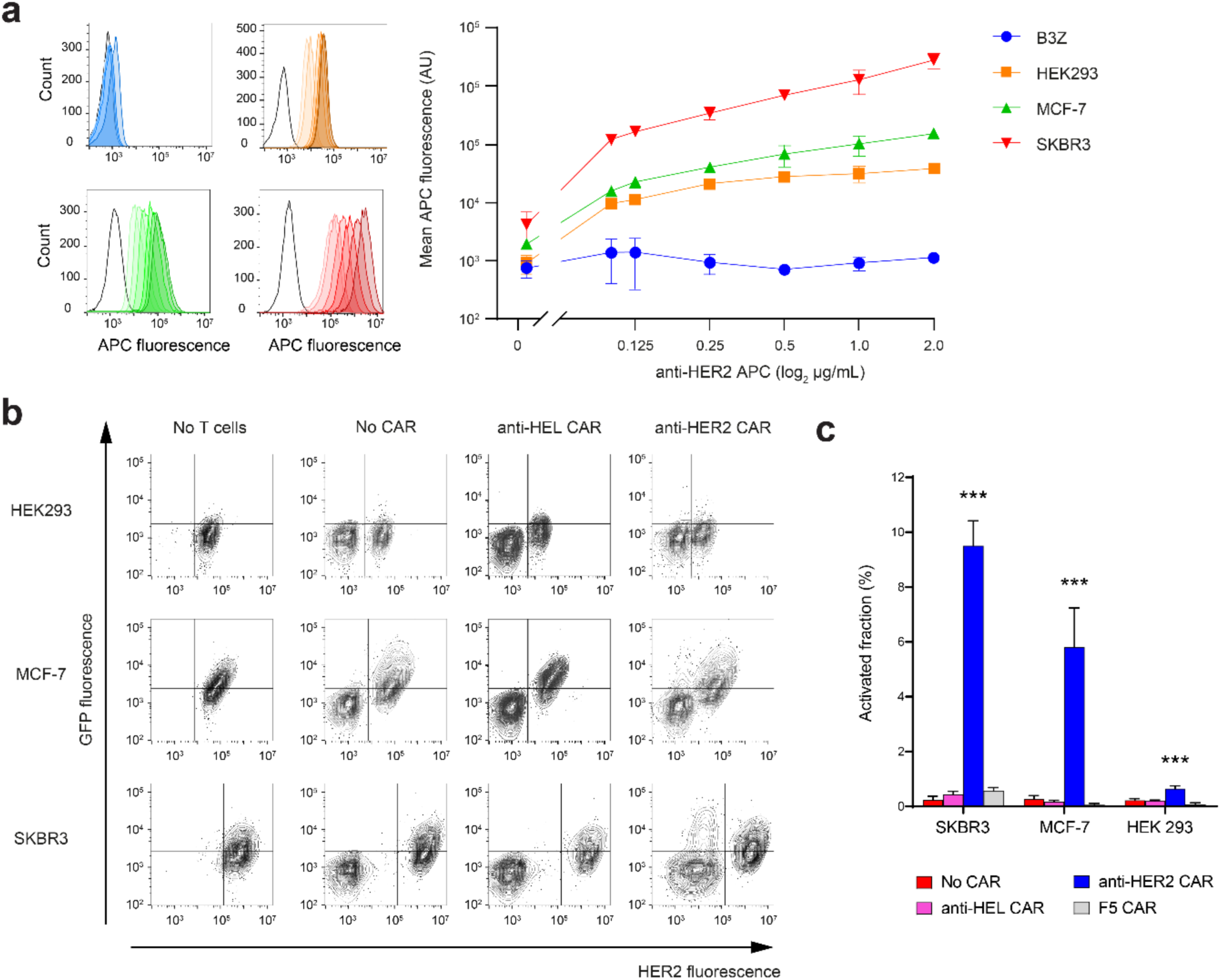
A CAR with a high affinity scFv shows a limited ability to discriminate between cell lines expressing different antigen levels. **a** Surface HER2 expression levels across the cell lines used in this study. Each cell line was stained with varying concentrations of anti-HER2 monoclonal antibody labeled with APC. Fluorescence was measured by flow cytometry and plotted in frequency histograms as follow: top left plot: B3Z; top right plot: HEK293; bottom left plot: MCF-7; bottom right plot: SKBR3. Mean fluorescence was plotted for all cell lines as a function of antibody concentration. SKBR3 shows the highest surface HER2 expression, followed by MCF-7, HEK293 and B3Z in descending order. **b** CAR T cells possessing a high affinity anti-HER2 scFv domain (4D5) were co-cultured with HER2-expressing cell lines. Following a 16-hour co-culture, cells were collected and flow cytometry was performed to measure HER2 and GFP expression. **c** Percentage of GFP-expressing T cells after co-culture with HER2-expressing cell lines. T cells expressing the Herceptin-derived anti-HER2 CAR showed a significant response to each tumour cell line compared to no CAR. Mean and S.E.M. were obtained from two independent experiments conducted with triplicates. For assessing significance, Dunnett’s multiple comparisons test was used with the following indicators: * P < 0.01, ** P < 0.001, *** P < 0.0001.

### Mutagenesis of a CAR and screening by antigen-mediated signaling or binding

The significant T cell activation encountered in the presence of both SKBR3 and MCF-7 cells suggested that the anti-HER2 CAR was not able to discriminate effectively between the antigen levels of each cell line. This is likely due to the very high affinity of the 4D5 scFv clone (equilibrium dissociation constant, *K_d_* ~ 0.1 nM ^26^). Using SKBR3 and MCF-7 cells to model tumour and healthy cells, respectively, we thus proceeded to use our T cell platform to engineer CAR variants that would retain activation in the presence of SKBR3 cells but show a reduced signaling response to MCF-7 cells. Our engineering strategy was based on the possibility of tuning the binding affinity of the scFv domain, such that a higher density of antigen molecules on the target cell’s surface is needed for successful T cell activation, thereby enabling greater cell-level selectivity. We hypothesized that our platform was suitable for this application as introducing mutations in the scFv domain will generate variants with different binding affinities and the IL-2-based signaling reporter could then be used to select for variants retaining responsiveness to HER2 on SKBR3 cells.

In order to generate a library of CAR variants with diverse HER2 antigen binding affinities, we performed deep mutational scanning (DMS) on the scFv domain. Specifically, we focused on the complementarity-determining region 3 of the variable heavy chain (CDRH3), which is a major determinant of binding specificity ^41^. The library was generated directly in T cells by genome editing, where Cas9 is used to integrate a pool of single-stranded oligodeoxynucleotide (ssODN) HDR templates to a pre-existing construct ^31, 42^. The HDR templates were based on a DMS design of a single-site saturation mutagenesis library, where degenerate codons (NNK; N = A, C, G T; K = G, T) are tiled across the 10 amino acids of the CDRH3 (Figure 4a). DMS libraries were screened and selected by FACS for cells with CAR surface expression based on Strep tag display. Next, deep sequencing was performed to assess the sequence landscape associated with CAR surface expression, demonstrating the expected diversity, with 190 out of possible 191 variants present and no apparent selection bias (Figure 4b).

**Figure 4:**
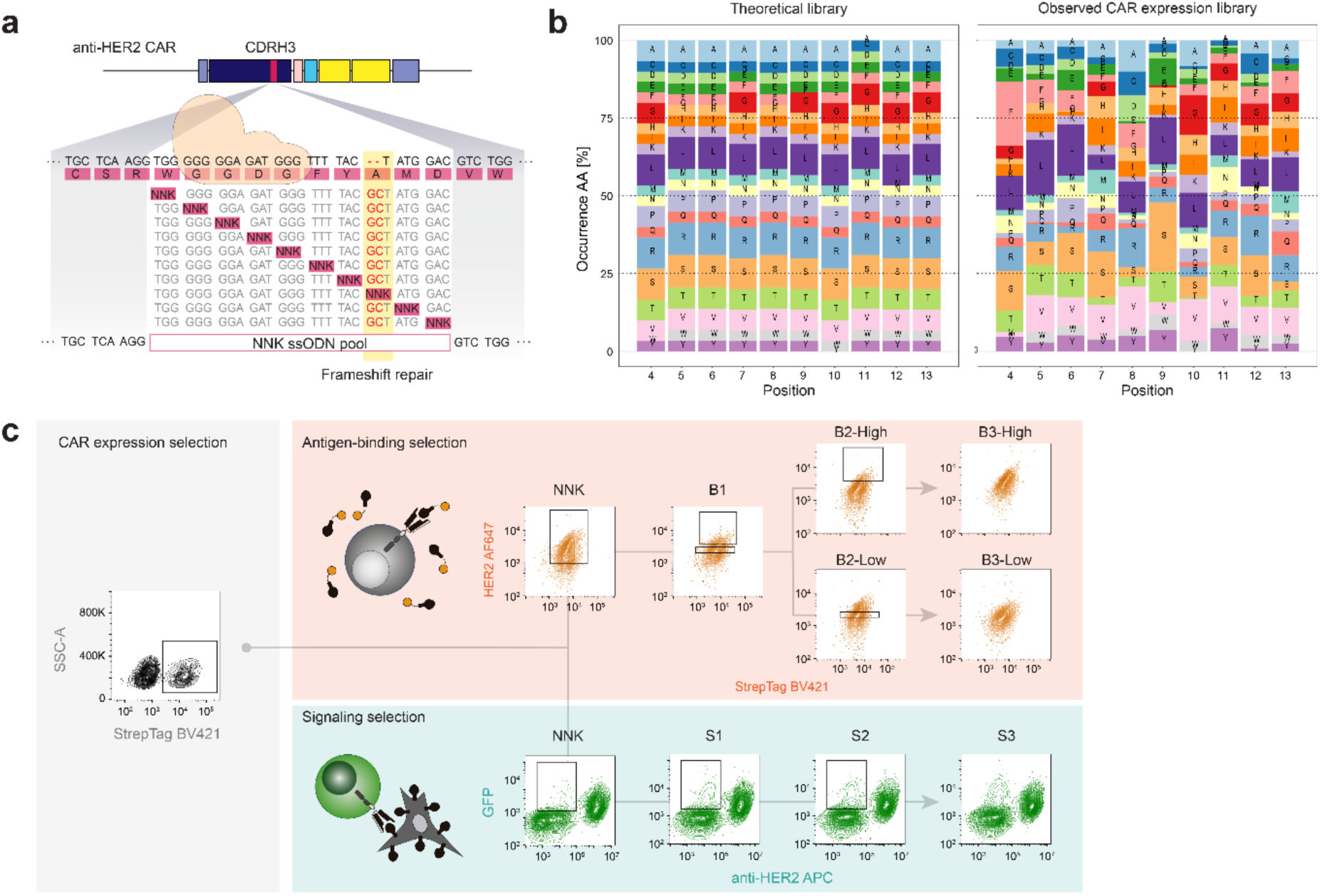
Generating a deep mutational scanning (DMS) library of CARs and selecting functional variants. **a** DMS of the CDRH3 region of the variable heavy chain of the CAR possessing the high affinity anti-HER2 scFv domain (4D5). The original anti-HER2 CAR was disrupted by the introduction of a frameshift-inducing deletion in the CDRH3. A set of degenerate ssODNs was then used to repair this deletion by CRISPR/Cas9 genome editing, while also substituting each position with all possible amino acids in a non-combinatorial fashion, for a total of 191 variants. Successful genome editing resulted in recovery of CAR surface expression, whenever possible. **b** Comparison of the theoretical (left) and observed (right) amino acid frequencies in the DMS library. Genomic DNA was extracted and the CDRH3 locus amplified and sequenced to measure the amino acid frequencies. The original 4D5 CDRH3 amino acids were not included in the histograms due to their overrepresentation. **c** FACS-based selection of CAR variants according to their antigen response or binding ability. Following transfection of the ssODN library, T cells expressing a CAR on their surface were selected based on Strep tag II staining (library “NNK”). This library was then used in two selection strategies in parallel. In one strategy based on antigen binding (“B”), a non-saturating concentration of soluble HER2 was used to select stronger (“High”) and weaker (“Low”) binders iteratively. In the second strategy based on signaling (“S”), co-culture with the high HER2-expressing cell line SKBR3 was followed by sorting according to GFP expression iteratively.

Next, we tested two approaches to select variants from the DMS library of CAR variants (Figure 4c). In one branch, selection was performed on the basis of binding to soluble HER2 antigen (“B” libraries). Iterative rounds of sorting were performed to select strong binders (B1, B2 High, B3 High) and weak binders (B1, B2 Low, B3 Low). In a second branch, CAR library variants were co-cultured with SKBR3 cells and IL-2 linked GFP expression was used to sort signaling responders (“S” libraries) in an iterative fashion (S1, S2, S3). The final libraries (B3-High, B3-Low and S3) were compared by their ability to respond to co-cultures with each of the HER2-expressing cell lines (Figure 5a and Supplementary Figure 3). The S3 library showed the highest ability to discriminate between the cell lines, largely due to unchanged signaling with SKBR3 and reduced signaling to MCF-7. Conversely, the B3 libraries showed reduced signaling to both cell lines compared to the initial CAR. These differences could not be accounted for by differences in CAR expression levels across libraries (Supplementary Figure 4).

**Figure 5:**
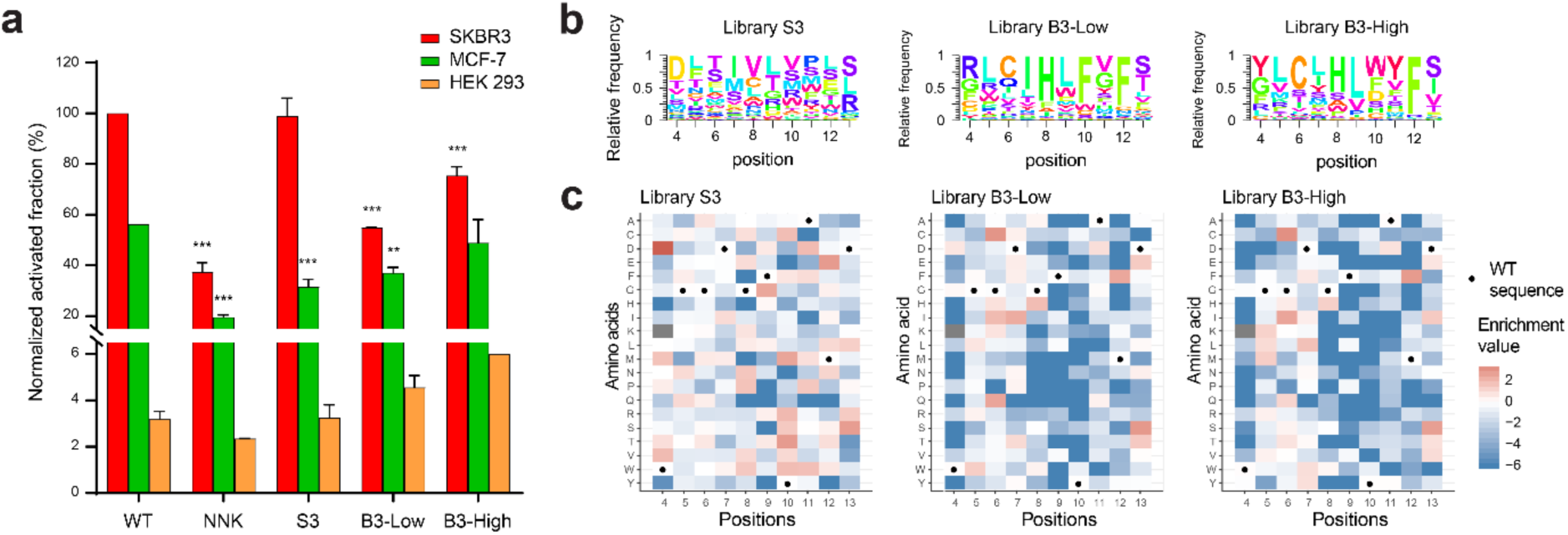
Deep sequencing of the libraries reveals the enrichment of different CAR variants based on the selection strategy. **a** Histogram of the response profiles of the wild-type anti-HER2 CAR T cells (WT), the initial (NNK) and the endpoint (S3, B3-Low, B3-High). The T cells were co-cultured overnight with HER2-expressing cell lines and the expression of GFP was measured by flow cytometry, revealing differences in the ability of libraries to discriminate between antigen levels compared to WT. Mean and S.E.M. were obtained from two independent experiments conducted with triplicates. For assessing significance, Dunnett’s multiple comparisons test was used with the following indicators: * P < 0.01, ** P < 0.001, *** P < 0.0001 **b** Sequence logo plots of the relative frequencies of the amino acid substitutions in the endpoint libraries. WT amino acids at each position are not shown. **c** Enrichment ratios for each CAR variant in the three endpoint libraries S3, B3-Low and B3-High in heat map form. Enrichment ratios were calculated as the ratio of the relative abundance of a sequence within an endpoint library over its relative abundance in the initial NNK library.

### Identification of CAR variants that are selectively activated based on tumour antigen surface expression

In order to identify single CAR variants showing an ability to discriminate across HER2 surface expression levels, we performed deep sequencing on the final binding and signaling libraries and compared the relative frequencies (Figure 5b) and the enrichment of single variants relative to the initial library (Figure 5c). The resulting enrichment heatmaps revealed more negative selection (in the form of lower enrichment scores) in libraries B3-High and B3-Low than for the S3 library, particularly at positions 8-12 of the CDRH3. Conversely, the S3 signaling library was less constrained across the entire region and deviated strongly from the other two, suggesting that there is a discordance between CAR antigen binding and signaling.

By using the deep sequencing data, we selected a set of variants from the S3 library based on high enrichment (Figure 6a and b). Additionally, we selected four variants from B3-Low showing more enrichment than in B3-High and three variants from B3-High. Cell lines of these CAR variants were then generated by Cas9-mediated HDR and subsequently co-cultured with either SKBR3 or MCF-7 and monitored for signaling activation by IL-2 linked GFP expression (Figure 6c and d, Supplementary Figure 5 and 6). Among them, four variants from the S3 library showed significant discrimination between the high-HER2 SKBR3 and low-HER2 MCF-7 cells when compared to the original CAR. Importantly, for variants M12E and Y10R, there was no reduction in responsiveness towards SKBR3. None of the variants from the binding libraries were able to discriminate based on HER2 surface expression, as they showed similar activation with both SKBR3 and MCF-7 cells, highlighting the difficulty of using antigen-binding as the only selection approach.

**Figure 6:**
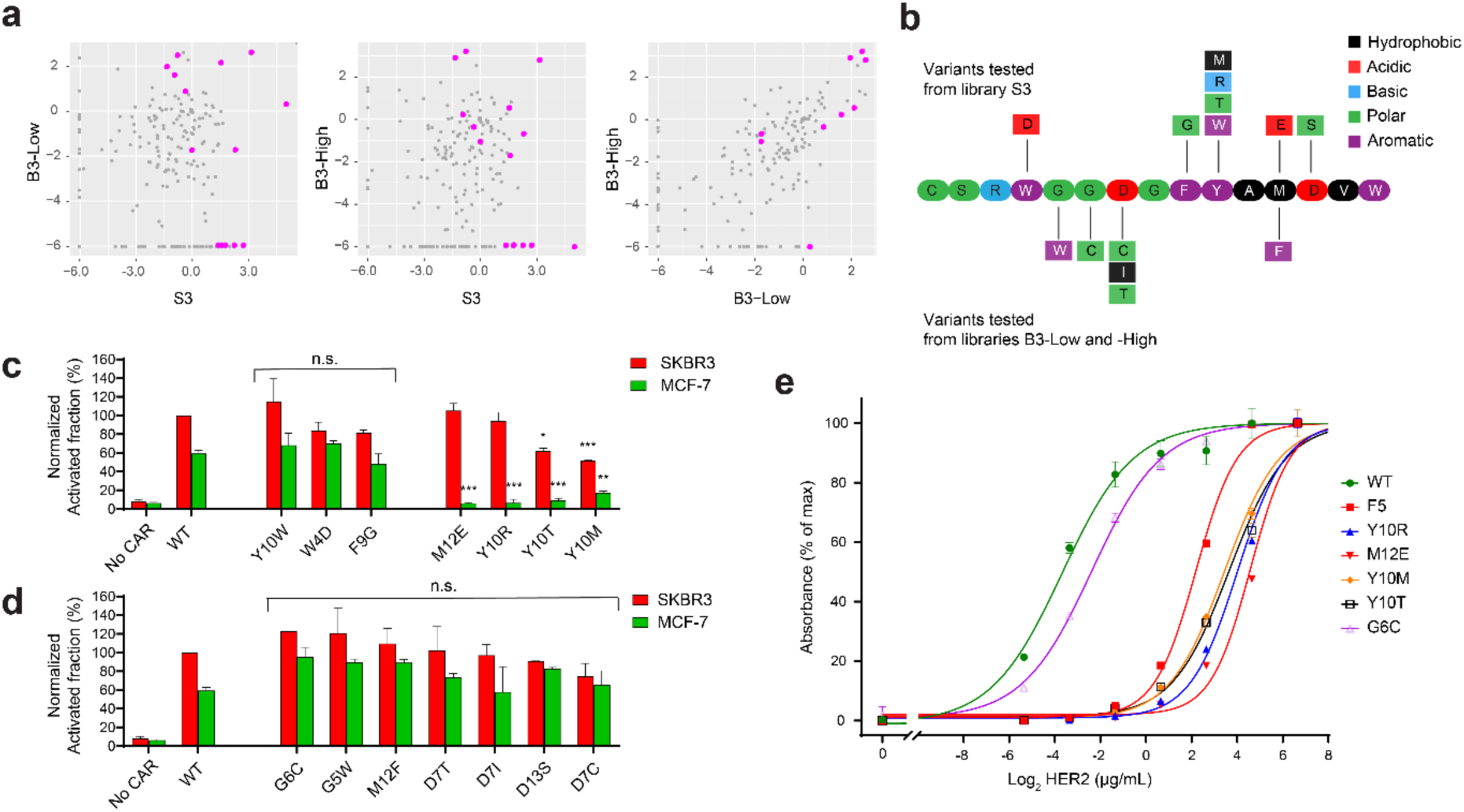
CAR variants enriched by signaling-based selection exhibit discriminative activated based on HER2 cell surface expression. **a** Scatter plot of the enrichment ratios for each variant across pairs of libraries. Variants selected for further study are highlighted in pink. The S3 library stands out as exhibiting little correlation to the two binding selection-based libraries. **b** Schematic of the CDRH3 region and the single-substitution variants individually selected based on their enrichment ratios. **c, d** Bar graphs of normalized activation (fraction of IL-2-linked GFP-positive cells) from CAR variants (selected in **a**) following co-culture with either SKBR3 (high HER2 expression) or MCF-7 (low HER2 expression). Four variants, selected based on signaling, (M12E, Y10R, Y10T and Y10M) respond significantly less to MCF-7 and of these, two (M12E and Y10R) show no significantly reduced signaling to SKBR3 compared to the wild-type (WT). No variants selected based on binding show a significant difference to the WT. Mean and S.E.M. were obtained from two independent experiments conducted with triplicates. For assessing significance, Dunnett’s multiple comparisons test was used with the following indicators: * P < 0.01, ** P < 0.001, *** P < 0.0001. **e** ELISA data of antigen-binding from CAR variants expressed in soluble scFv form. The variable heavy and light chains of the CAR variants were expressed in bacteria with a His tag. Following immobilization on a plate with a monoclonal anti-His tag antibody, binding to varying concentrations of biotinylated HER2 was detected with the addition of Streptavidin-HRP.

Finally, we aimed to quantitatively understand the impact of antigen-binding of our CAR variants that showed discriminative activation based on HER2 surface expression. We expressed the binding domain of the CARs in soluble scFv form and measured their binding to HER2 antigen by ELISA (Figure 6e). This assay revealed strong decreases in binding for all of the discriminating variants. By comparison, the variant G6C, obtained by binding selection, showed only a modest reduction in binding relative to the initial 4D5 scFv. Furthermore, we found that the variant F5, which was largely unresponsive to SKBR3 (Figure 3c), displayed a binding curve similar to that of the discriminating variants, highlighting a divergence between cell responsiveness and antigen-binding. Taken together, this suggests that signaling-based selection is a superior strategy to binding-based selection for identifying CAR T cell variants that can discriminate based on tumour antigen surface expression.

## DISCUSSION

Through a series of genome editing steps, we have engineered a CAR T cell display platform for screening based on signaling based-activation or antigen binding. We have demonstrated its value for affinity tuning, a simple and yet effective method to reduce the likelihood of on-target, off-tumour effects. This approach relies on general principles of genome editing and directed evolution, and has the potential to eliminate instances of toxicity early in the development pipeline of CAR-based immunotherapies. The benefits of decoupling receptor affinity and T cell activation were first shown by Chmielewski *et al.*, which in a study found that CARs with *K_d_* affinity values above 10^-8^ M responded strongly to high HER2-expressing cell lines in vitro, but were not activated in the presence of low HER2-expressing lines, unlike their higher affinity counterparts ^43^. This result encapsulates a paradox of affinity in the context of cell-mediated cytotoxicity: while high affinity theoretically implies exquisite specificity at the epitope level, specificity at the cell/tissue level can in fact be impaired due to an inability to discriminate between antigen levels. Furthermore, there may be additional benefits to reducing CAR affinity, such as greater cell expansion and longer persistence ^44^. These observations are likely a factor for the characteristic low binding affinity of endogenous TCRs relative to that of B cell receptors and antibodies ^45^. In T cell maturation, central tolerance mechanisms ensure that high-affinity self-reactive T cells are removed from the immune repertoire. For CAR engineering, a process must be devised with an equivalent outcome to avoid targeting cells expressing normal levels of oncogenes.

Our CAR display platform closes existing gaps in current methods of tuning target selectivity. Previous examples have relied on utilizing low affinity antibodies to rationally design CARs with safer target selectivity ^24, 25, 26, 27, 28, 29^. While our platform can also be used to rapidly assess the functionality of rationally designed variants, its true strength lies in its ability to accommodate library generation and functional screening. Given the uncertainty surrounding the threshold of affinity required for CAR triggering, optimizing CAR safety benefits greatly from a readout for cell signaling. Here, the IL2-GFP reporter acted as a “guardrail”, preventing selected variants from “falling” into non-functionality. This is especially important since the activation threshold varies across antigens and epitopes ^26, 43^, and may vary across CAR signaling components (e.g., CD28 vs. 4-1BB co-signaling domain) ^46^. Differences in epitopes may in fact explain why no activity was observed with the CAR scFv variant F5, which was derived from a phage display library screened for triggering the intracellular uptake of bound HER2 ^39^. Epitope mapping suggests that it targets the membrane-distal domain 1 of HER2, while the 4D5 antibody targets the membrane-proximal domain 4 ^39, 47^. It is not immediately clear why this difference affects CAR triggering, despite the relative binding of F5 being seemingly comparable to that of our variants which maintain signaling. Until such questions are resolved, a display platform such as ours that enables the detection of signaling activation ensures that no prior knowledge regarding the strength of the affinity or the targeted epitope is required for optimizing a CAR.

Engineering a CAR for selectivity based on tumour antigen surface expression relies on the important notion that antigen-binding affinity and antigen-induced signaling strength can diverge. This is most apparent for CARs with high binding affinities, where a “ceiling” is encountered above which activation does not improve further ^23, 43^. This ceiling implies that reducing CAR affinity does not necessarily lead to a decline in responsiveness to a target tumour, and so signaling is a more “relaxed” constraint than binding. This was evident as we attempted to also select low affinity variants by antigen binding selection, but this was not successful on its own in identifying variants that could discriminate on the basis of tumour antigen surface expression. A possible reason for this is that our selection (gating) strategy was too conservative and did not sufficiently deviate from the B3-High strategy. The difficulty of selecting an appropriate gating strategy further supports the use of signaling-based selection which was trivial by contrast, merely requiring that all GFP-positive clones be selected. Furthermore, although our end goal was to improve discrimination between two cell lines, SKBR3 and MCF-7, it is notable that the latter played no part in our selection strategies. As the difference in antigen levels between the two cell lines was relatively large, no negative selection step was required. The binding libraries might have benefited from negative selection, but this was constrained by the low sensitivity of fluorescence-based binding detection, which made it difficult to distinguish weak-binders from non-binders. This became especially apparent once we had obtained discriminating variants by signaling selection, as these showed no detectable binding to soluble antigen by flow cytometry, despite co-culture and ELISA assays confirming their function. Signaling pathways can amplify signals that are mistakenly deemed too weak to trigger a response, even when the reporter includes significant signal dampening through negative regulation ^48^. The endogenous IL-2 promoter offers a suitable sensitivity as a reporter of CAR T cell activation for screening purposes.

Our affinity tuning approach also benefited from the combination of DMS, which can cover a large sequence space in an unbiased way, and from deep sequencing, which revealed the substitutions that became enriched as a result of different selection approaches. Performing a single-site saturation mutagenesis scan can avoid shifting the target epitope too far from what has been deemed a good target site, saving us from “reinventing the wheel” by screening novel variable heavy and light chain combinations. As the majority of random mutations do not improve a protein’s function, we hypothesized that a combination of random mutagenesis and relaxed selection constraint would broaden and left-skew the distribution of binding affinities. After a single-mutation scan, our experiment resulted in a moderate improvement in antigen-level discrimination across the whole library level solely through the signaling-based selection strategy, validating this approach. However, the binding selection strategies also proved essential for identifying the specific mutations responsible for this phenotype. The enrichment of our best variants, M12E and Y10R, did not lead them to dominate the S3 library, since the strong HER2 binding variants were not depleted by the relaxed signaling-based constraints. Rather, the depletion of M12E and Y10R in the B3 libraries was key for highlighting their tuned affinity. In fact, the Y10 position showed strong negative selection in these libraries, revealing its value for discriminative activation. The use of DMS and parallel selection strategies thus appears to be a promising method for gaining insight on key residues that can be altered without abrogating the antigen-specific signaling response.

As the CAR immunotherapy field shifts towards addressing solid tumours, an increasingly diverse array of antigens requires targeting, and so more instances of off-tumour toxicity will be encountered. This does not have to lead to the exclusion of valid targets and abandoning the wide range of high-affinity antibodies that have been discovered to date. Rather, it is likely that many of these are amenable to tuning CAR selectivity at little or no cost to the maximal signaling response, provided that the target antigen is overexpressed on malignant cells. Our CAR display platform facilitates this process by enabling library generation and high-throughput functional screening. The simplicity of this method may present notable advantages versus more complex approaches (e.g., combinatorial logic gates and soluble molecules) that have been put forward to deal with off-tumour toxicity effects. Alternatively, engineering CAR selectivity based on antigen affinity could be combined with such methods. For instance, multi-specific CAR T cells have recently been developed that integrate the recognition of two targets ^49, 50, 51^. In such a system, it is crucial that no single binding domain can trigger a cytotoxic response on its own ^16^. Such constructs could potentially be balanced by tuning CAR antigen binding as described here. Functional display platforms can thus accelerate the development of increasingly sophisticated cell-based immunotherapy treatments that are both effective and safe.

## METHODS

### Cell culture

B3Z cells were cultured in Iscove’s Modified Dulbecco’s Medium (IMDM) with Glutamax; SKBR3, MCF-7 and HEK293 cells were cultured in Dulbecco’s Modified Eagle Medium (DMEM) (Gibco). Culture media were supplemented with 10% fetal bovine serum, 1% Penicillin-Streptomycin (Gibco) and 100 µg/mL Normocin (Invivogen). For passages and experiments, adherent cell lines were detached using TrypLE™ Express (Thermo Fischer) at 37°C.

### Genome editing and sequencing

Genomic modifications of B3Z cells were performed by CRISPR/Cas9 genome editing. Guide RNA (gRNA) was assembled as a crRNA (Supplementary Table 2) and tracrRNA duplex according to the manufacturer’s instructions (IDT). For cell lines without endogenous Cas9 expression, a ribonucleoprotein (RNP) was assembled by incubating the duplex RNA and *Streptococcus pyogenes* Cas9-V3 (IDT) at room temperature for 20 min. For amplimer HDR, a double-stranded DNA repair template was generated by PCR with flanking homology arms of ~700 bp in length; 5 µg of purified product was used for the transfection. For ssODN HDR, 500 pmol ultramer (IDT) was used. Transfections (electroporations) were carried out using a Lonza 4D-Nucleofector according to recommended protocols. Briefly, 500,000 B3Z cells were collected and resuspended in SF buffer. The RNP/DNA mixture was added to the cells in a 1:10 ratio for a total volume of 100 µL. Electroporations were performed in Nucleocuvettes with the program CA-138. Cells were then diluted in 600 µL warm medium. Assays or sorting were performed at least four days later. To confirm genome editing, genomic DNA was extracted from at least 10^4^ harvested cells using the QuickExtract protocol (Lucigen). The target locus was then amplified by PCR with forward and reverse primers where at least one primer annealed outside the integration site (Supplementary Table 1). Sanger sequencing was used to confirm correct integration of HDR templates.

### Flow cytometry and cell labeling

The expression of surface markers or of the genomically-integrated green fluorescence protein (GFP) were assessed by flow cytometry. Cells were first washed in Dulbecco’s PBS (DPBS) prior to labeling. For CAR detection, labeling with 1:200 biotinylated anti-Strep tag antibody (GenScript) was followed by 1:500 BrilliantViolet 421 conjugated with Streptavidin (Biolegend). To assess HER2 binding, cells were incubated with 2.5 µg/mL soluble HER2 antigen (Merck) and then with 1:250 APC-labeled anti-human HER2 antibody (Biolegend). The same antibody and concentrations were used to measure surface HER2 expression on cell lines SKBR3, MCF-7 and HEK293. For TCR expression, cells were stained with 1:200 APC-labeled anti-mouse Vβ5.1/5.2 antibody (Biolegend). For CD3ε expression, cells were stained with 1:200 APC-labeled anti-mouse CD3ε antibody (Biolegend). Cells were kept in DPBS on ice until analytical flow cytometry or FACS. Flow cytometry data were analysed by Flowjo.

### Co-culture assays and sorting

CAR-expressing B3Z cells were co-cultured in a 1:1 ratio with HER2-expressing cells (SKBR3, MCF-7 or HEK293) in B3Z medium for 16 hours. All cells were then collected, washed in DPBS and stained for flow cytometry analysis or sorting. For assays, 2.5×10^4^ B3Z cells were used, while 3×10^6^ cells were used for sorting.

### Deep mutational scanning

The DMS CAR library was generated by CRISPR-Cas9 genome editing of the 4D5 CDRH3 with a pool of single-stranded DNA HDR mutant repair templates, as done previously ^31^. Briefly, gRNA targeting the CDRH3 of the anti-HER2 CAR in the CAR T cell platform was used to obtain a monoclonal T cell line with a frameshift deletion abrogating CAR expression. This deletion was then repaired with a pool of HDR templates encoding single amino acid substitutions of the CDRH3. The pool was designed with tiled degenerate codons such that all possible amino acid substitutions were represented along 10 positions of the CDRH3. The HDR templates were transfected in the non-CAR expressing T cells, along with gRNA targeting the CDRH3 deletion. To identify enriched CAR variants post-selection, deep sequencing of the libraries was done by a nested PCR strategy to amplify a 401 bp fragment from genomic DNA, which included the CDRH3. A first PCR was performed to amplify the entire CAR transgene; the product was then used as a template for amplification of the smaller CDRH3-containing region. Second, the product was used in a PCR with primers CDRH3_seq_F and CDRH3_seq_R (Supplementary Table 1). The resulting fragments were purified and sequenced by GENEWIZ (Leipzig, Germany). Only sequences with a complete CDR3 harboring a single amino acid substitution (resulting from DMS) were considered in the analysis. The relative abundance of each variant was extracted and used to calculate enrichment ratios with respect to the initial CAR library (“NNK” in the text).

### Recombinant expression and measurement of scFv binding to antigen

Soluble scFv proteins were recombinantly expressed: the coding sequences were cloned in a bacterial pET28 expression vector with a C-terminal His tag and transformed into BL21-DE3 competent *E. coli* cells (NEB). Cells were grown in LB medium at 37°C to an optical density at 600 nm (OD_600_) of 0.6 and protein expression was induced with 1 nM IPTG for 24 hours at 20°C. Cells were then harvested by centrifugation, and proteins were recovered from the periplasm by sonication. TALON metal affinity chromatography was used for purification as previously described ^52^. ELISA assays were done with material and reagents from Thermo Fisher Scientific. For this, soluble HER2 antigen was first biotinylated with the EZ-Link™ NHS-PEG4-Biotin reagent for eventual detection. Nunc Maxisorp 96-well plates were coated with 2 µg/mL mouse IgG2b monoclonal 6x-His tag antibody (HIS.H8) and the subsequent blocking was done with ELISA/ELISPOT diluent. The His-tagged scFv proteins were allowed to bind at a concentration of 2 µg/mL, followed by varying concentrations of biotinylated HER2 antigen, for 2 hours each. Detection was done with tetramethylbenzidine (TMB) substrate solution and 1:1000 Ultra Streptavidin-HRP. The enzymatic reaction was stopped with 0.16 M sulfuric acid. Readings were done with a Infinite Pro M200 plate reader (Tecan) at 450 nm with subtraction at 570 nm.

## AUTHOR CONTRIBUTIONS

R.B.D.R and S.T.R. designed the study; R.B.D.R., R.C.R., S.F., D.E. and R.V.-L. performed experiments; R.B.D.R., R.C.R. and S.T.R. discussed results. R.B.D.R., R.C.R. and S.T.R. wrote the manuscript with input and commentaries from all authors.

## ACKNOWLEDGMENTS

We acknowledge the ETH Zurich D-BSSE Single Cell Unit for excellent support and assistance. This work was supported by a ETH Zurich Post-doctoral Fellowship (to R.B.D.R.), a Personalized Health and Related Technologies Post-doctoral Fellowship (to R.V.-L.) and a NCCR Molecular Systems Engineering (to S.T.R.)

## DATA AVAILABILITY

The raw FASTQ files from deep sequencing that support the findings of this study will be deposited (following peer-review and publication) in the Sequence Read Archive (SRA) with the primary accession code(s) <code(s) (https://www.ncbi.nlm.nih.gov/sra)>. Additional data that support the findings of this study are available from the corresponding author upon reasonable request.

