## Supplementary Material for "A functional screening platform for engineering chimeric antigen receptors with reduced on-target, off-tumour activation"

**SUPPLEMENTARY INFORMATION**

SUPPLEMENTARY FIGURES 1 – 6

SUPPLEMENTARY TABLES 1 and 2

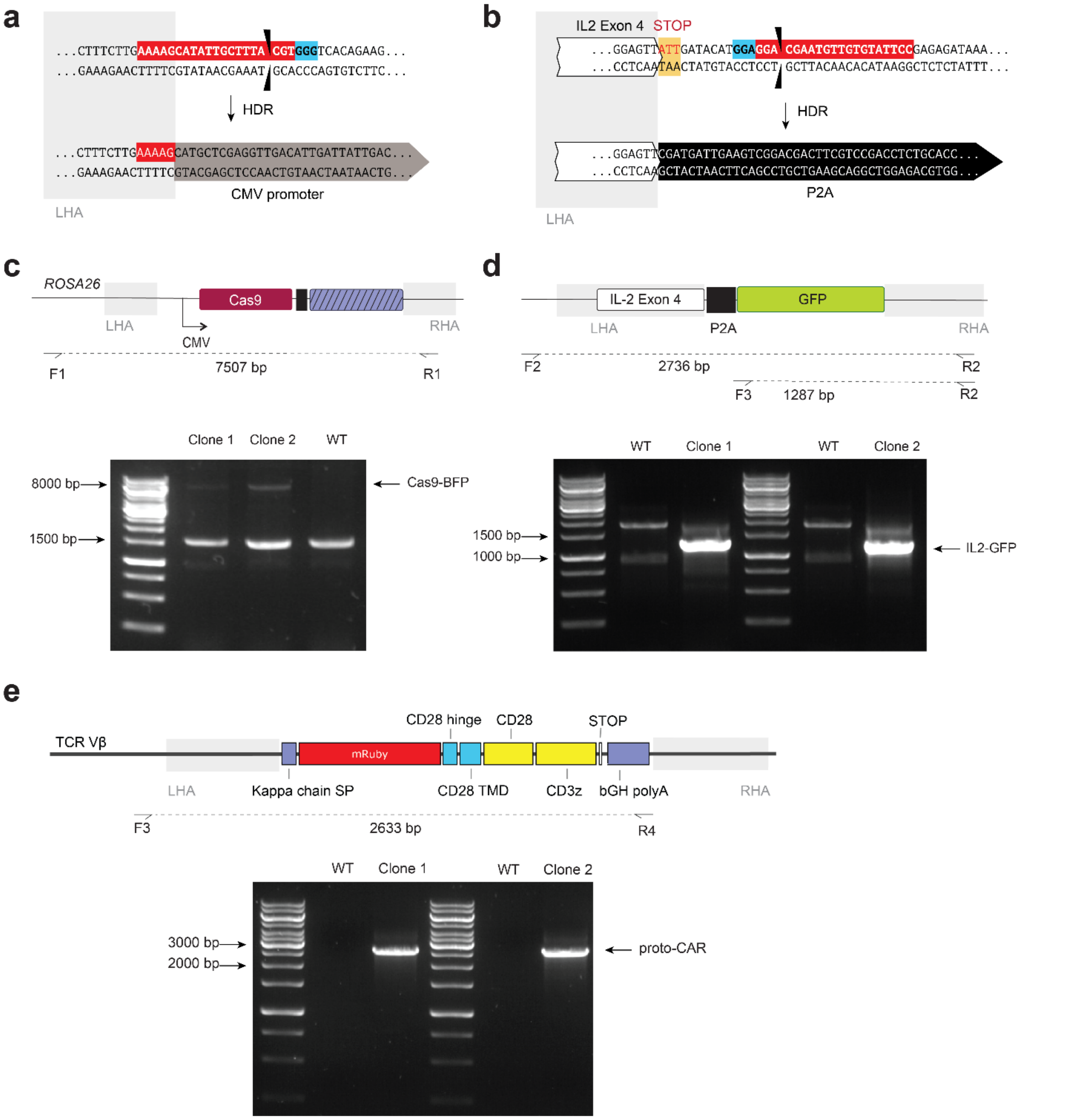


**Supplementary Figure 1: Genomic integration strategy and PCR confirmations for engineering the CAR T cell platform.** **a** CRISPR-Cas9 cut site in the *ROSA26* locus and post-HDR sequence of the integrated Cas9-BFP expression cassette. **b** CRISPR-Cas9 cut site in the IL-2 locus and post-HDR sequence of the integrated GFP ORF. **c-e** PCR amplification strategy to confirm the genomic integration of the Cas9-BFP cassette, GFP ORF or proto-CAR expression cassette. DNA gels show the sequenced amplimers.

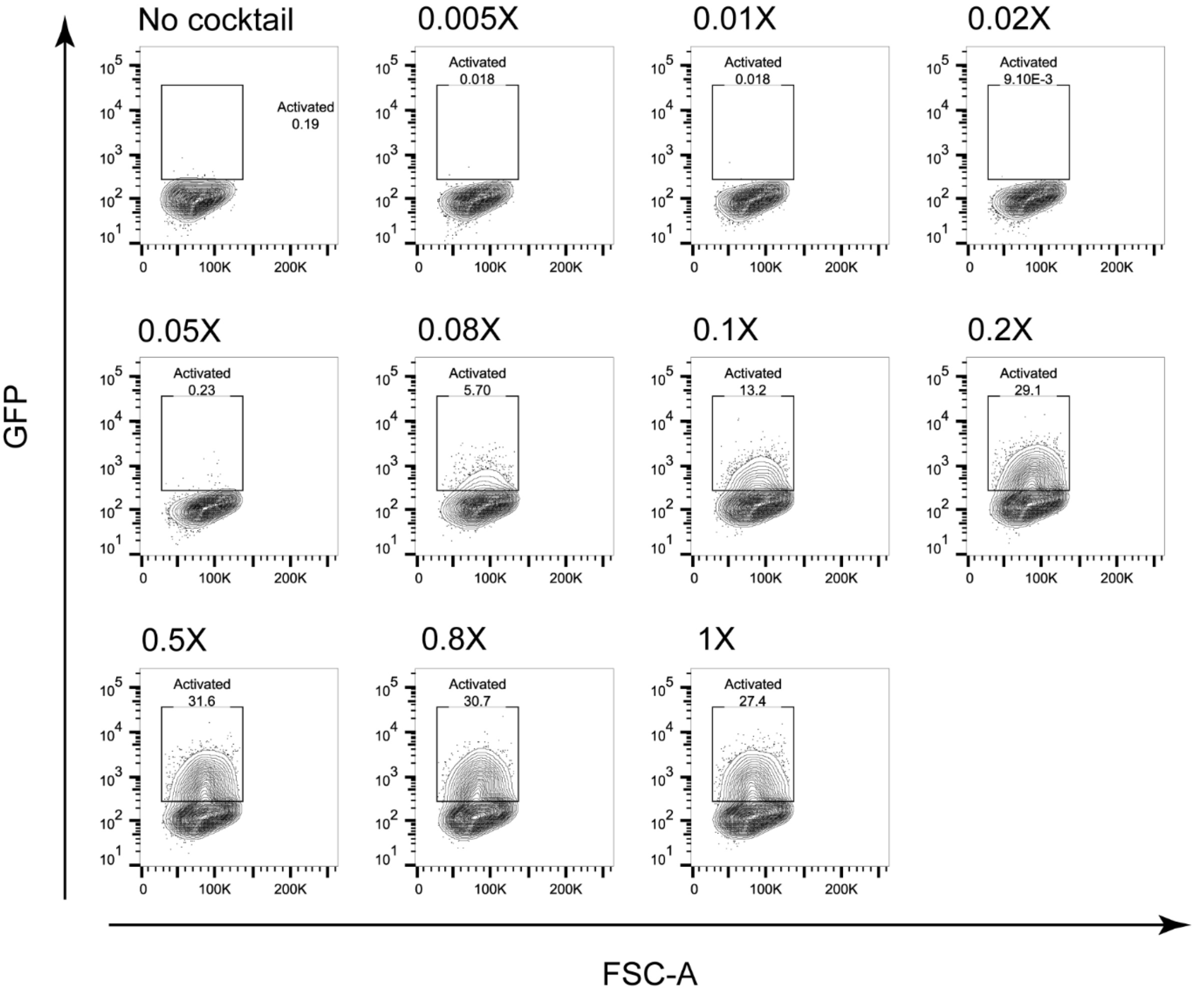


**Supplementary Figure 2: Representative flow cytometry plots of GFP expression in engineered T cells following PMA/ionomycin stimulation.** Following overnight incubation with varying concentrations of the cell stimulation cocktail PMA/ionomycin (1X: 2 μL/mL), cells were collected and assayed. The fraction of GFP-positive cells was used to plot the dose-response curve of Fig. 1**d**.

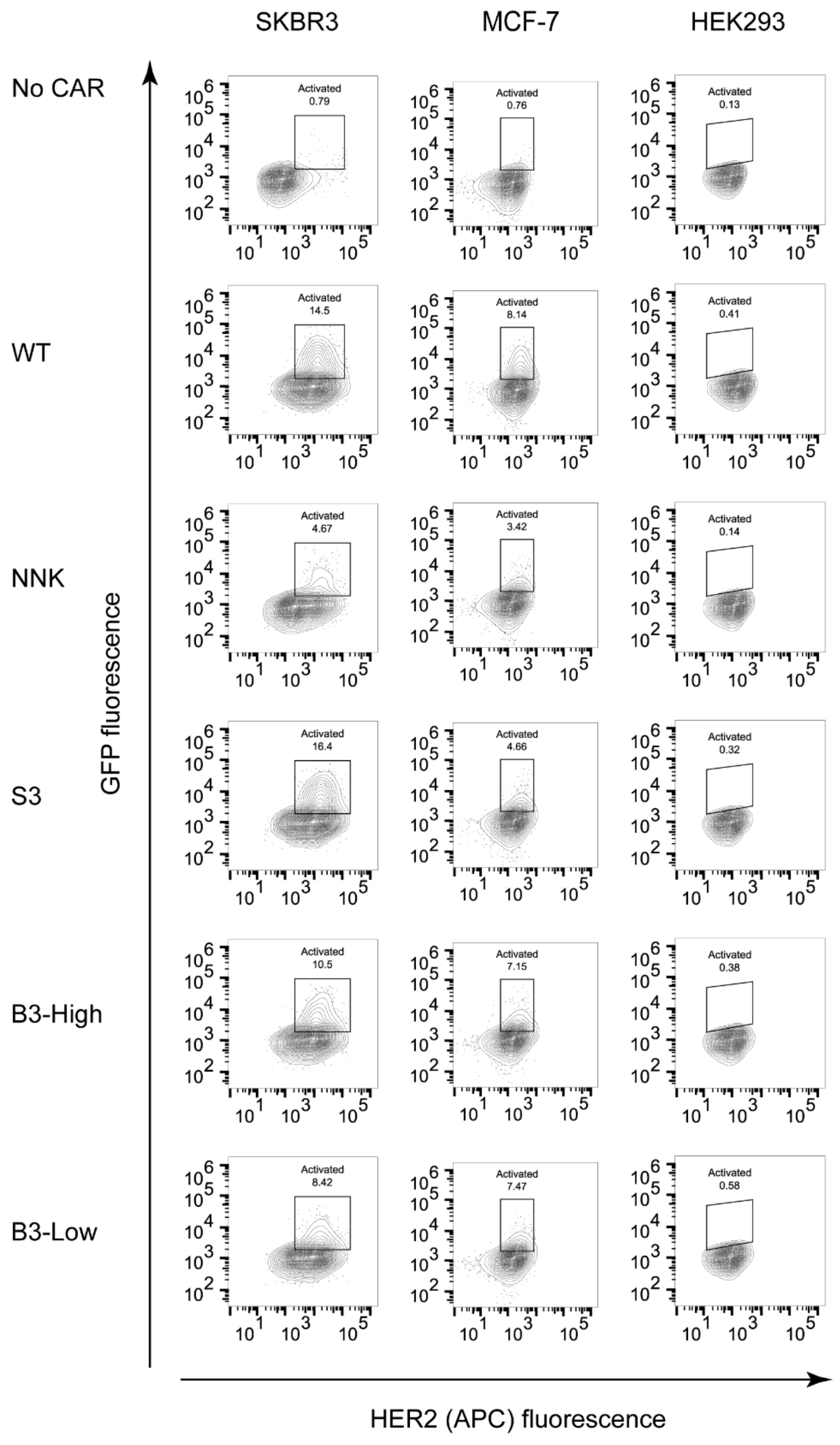


**Supplementary Figure 3: Representative flow cytometry plots of GFP expression in post-selection CAR T cell libraries following co-culture with HER2-expressing cell lines.** CAR T cell libraries were incubated overnight in co-culture with SKBR3, MCF-7 or HEK293. Cells were collected and labeled with monoclonal anti-HER2 antibody (APC-conjugated) to distinguish CAR T cells from target cells on the basis of HER2 cell surface expression. The fraction of GFP-positive cells was used to plot the histogram of Fig. 4**a**.

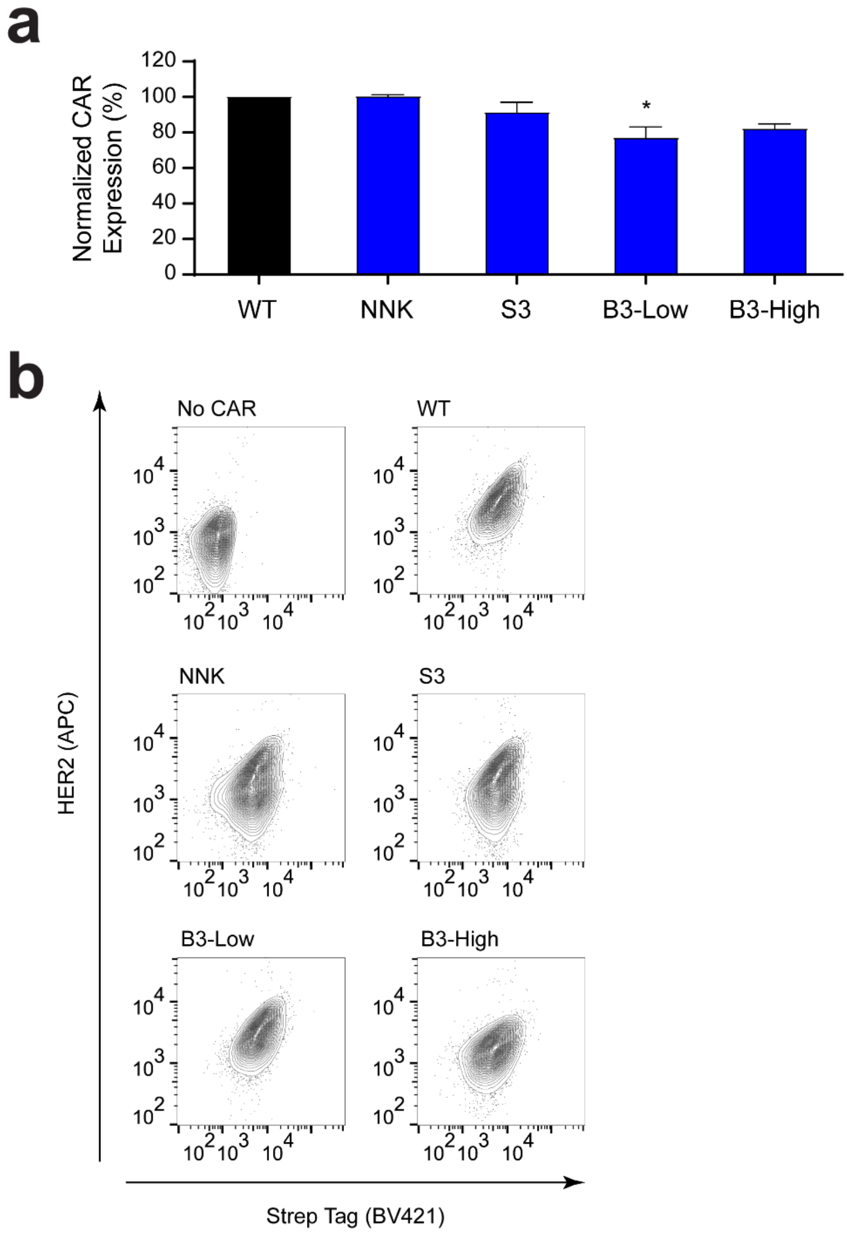


**Supplementary Figure 4: CAR cell surface expression shows little difference across variant libraries. a** Bar plot of CAR surface levels from CAR T cell platform libraries following selection. Mean fluorescence for each was obtained from two independent experiments normalized to the WT of that experiment after subtracting background fluorescence. For assessing significance, libraries were compared to the WT and Dunnett's multiple comparisons test was used with the following indicators: * P < 0001, ** P < 0.001, *** P < 0.0001. **b** Representative flow cytometry plots of CAR T cells used to generate the bar plot in **a**. Cells were labeled with a monoclonal anti-Strep tag (BV421-conjugated) and a combination of soluble HER2 antigen and monoclonal anti-HER2 antibody (APC-conjugated).

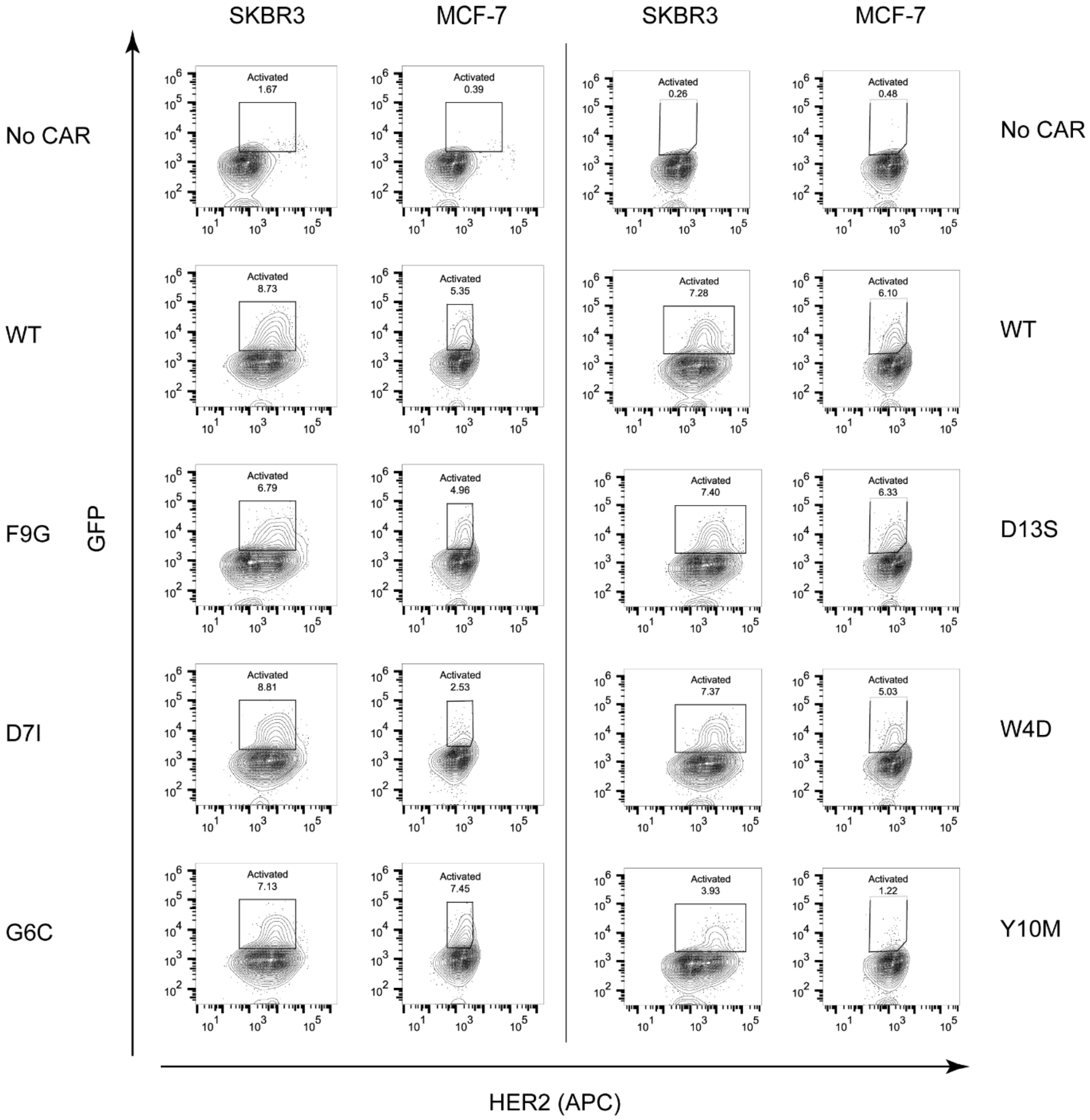


**Supplementary Figure 5: Representative flow cytometry plots of GFP expression in CAR T cell variants following co-culture with HER2-expressing cell lines.** Selected CAR T cell variants were incubated overnight in co-culture with SKBR3 or MCF-7 cells. Cells were collected and labeled with monoclonal anti-HER2 antibody (APC-conjugated) to distinguish CAR T cells from target cells on the basis of HER2 surface expression. The fraction of GFP-positive cells was used to plot the bar plot of Fig. 5**c** and **d**.

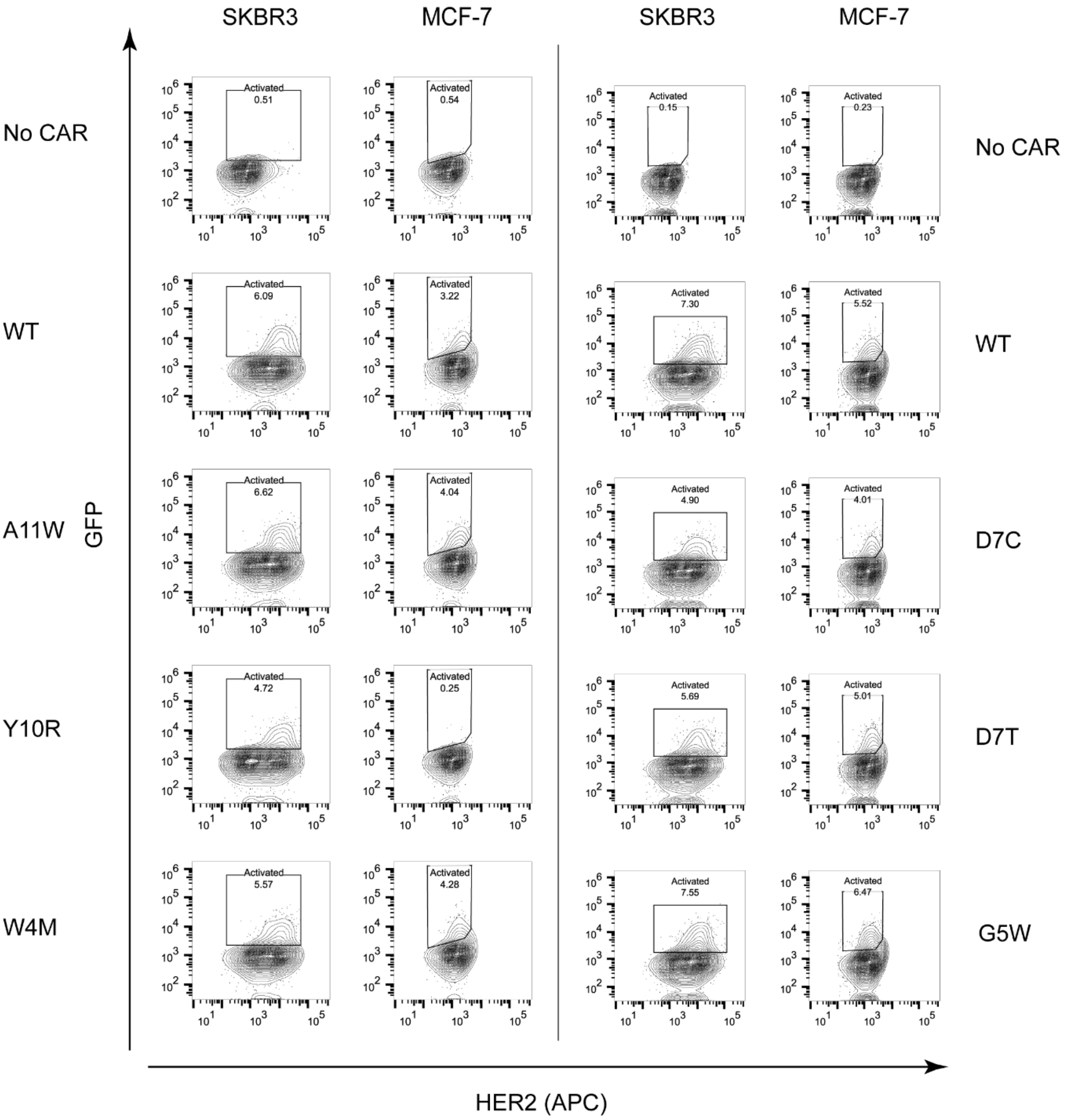


**Supplementary Figure 5 (continued)**

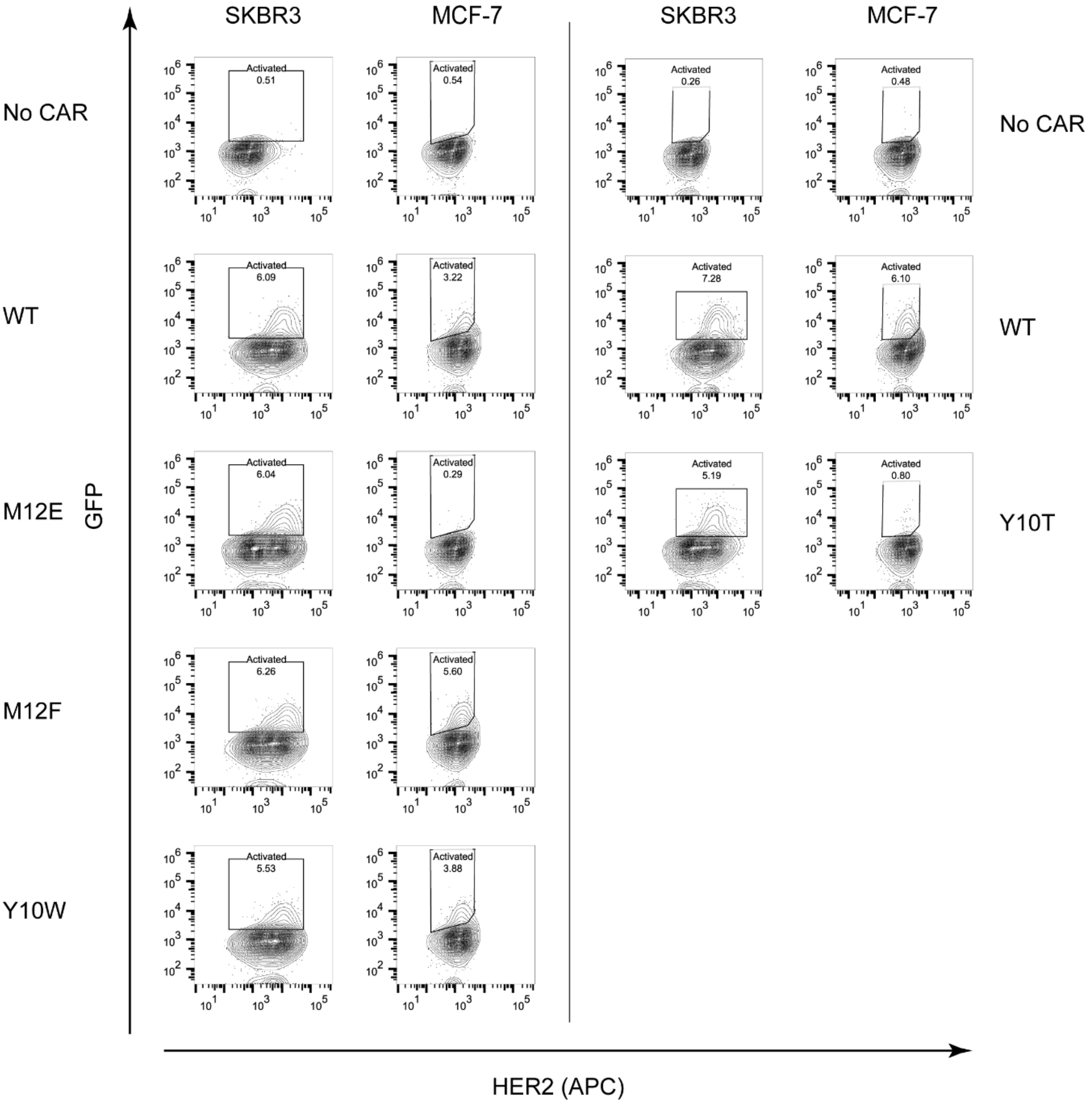


**Supplementary Figure 5 (continued)**

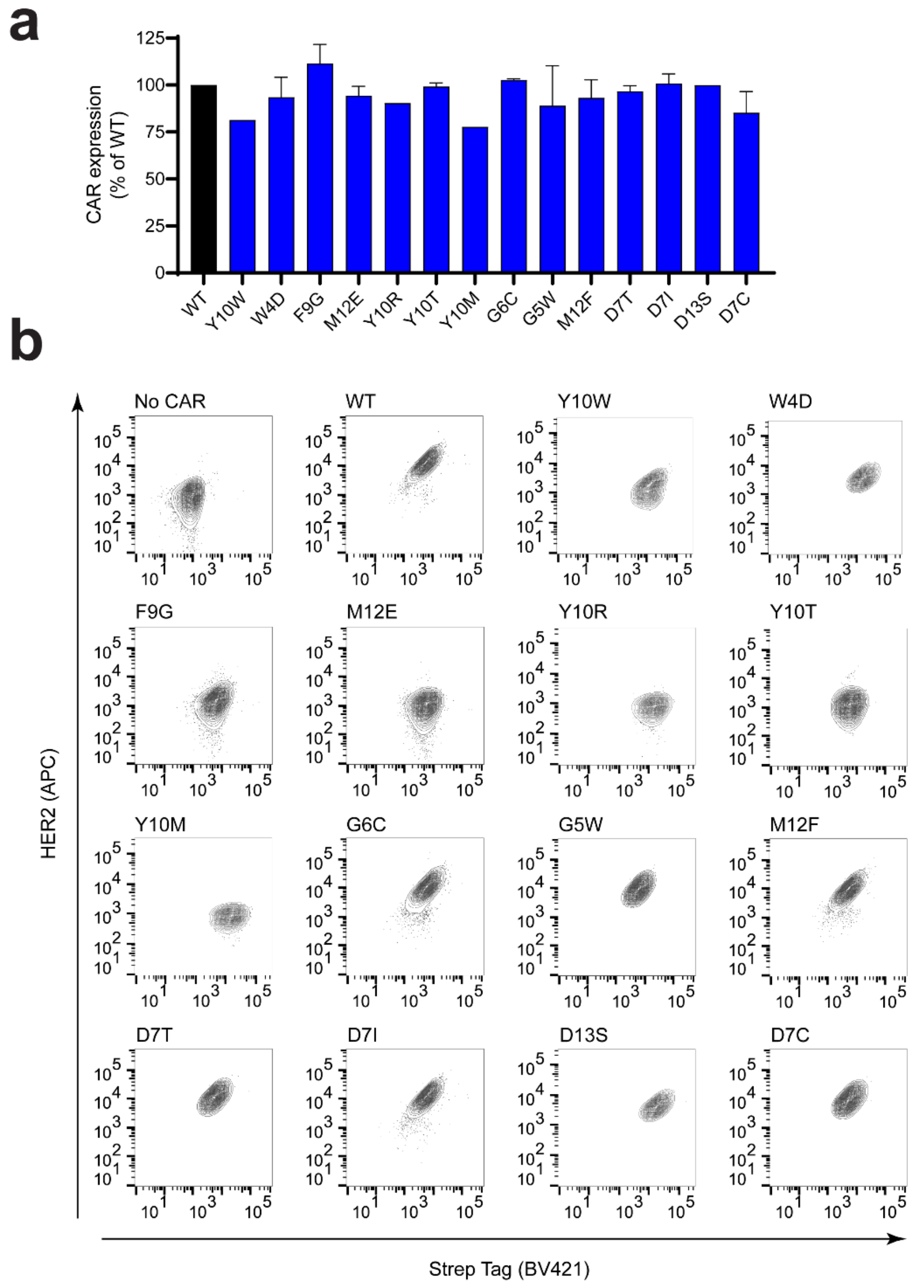


**Supplementary Figure 6: CAR surface expression shows little difference across variants. a** Bar plot of CAR surface expression from selected CAR T cell variants. Mean fluorescence for each was obtained from two independent experiments normalized to the WT of that experiment after subtracting background fluorescence. For assessing significance, variants were compared to the WT and Dunnett's multiple comparisons test was used with the following indicators: * $P < 0001$, ** $P < 0.001$, *** $P < 0.0001$. **b** Representative flow cytometry plots of CAR T cells used to generate the bar plot in **a**. Cells were labeled with a monoclonal anti-Strep tag (BV421-conjugated) and a combination of soluble HER2 antigen and monoclonal anti-HER2 antibody (APC-conjugated).

SUPPLEMENTARY TABLES

Supplementary Table 1: Oligonucleotides used in this study

| **Name** | **Sequence (5’ to 3’)** | **Purpose** |
|---|---|---|
| F1 | CTGTATTAGTTGAAGGAAAGTGT | To amplify Cas9-BFP in the *ROSA26* locus for sequencing |
| R1 | TTTACTGAACAAATGAACTACTTTTT | |
| F2 | TTGCCTTTGCCAAAATAAGATTTTA | To amplify IL2-GFP in the IL-2 locus for sequencing |
| R2 | AAGAGGGTGTGGGCTAGTCCTTTCC | |
| F3 | ATGGTGAGCAAGGGCGAGG | |
| F4 | GAGCCCCTTTTTTTTTTGGTCTG | To amplify the CAR in the TCR Vβ locus |
| R3 | CTTACCCATGCCTGCTATTCTCTTCCCAATC | |
| F5 | GGTCAGACAAGCTCCCGGAAAAGGA | To amplify the CDRH3 of the anti-HER2 CAR for sequencing |
| R4 | TTCCATTGCTCCTCTCGTTGTCTAGG | |

Supplementary Table 2: crRNA sequences used in this study

| **Genomic target** | **crRNA sequence (5’ to 3’)** |
|---|---|
| *ROSA26* | AAAAGCAUAUUGCUUUACGU |
| BFP | CACUUCAAGUGCACAUCCGA |
| IL-2 | GGAAUACACAACAUUCGUCC |
| mRuby2 | GUUUACCACGUCCAAGUCAG |
| 4D5 CDRH3 | GGGAGAUGGGUUUUACGCUA |
| 4D5 CDRH3 (with deletion) | GUUUUACUAUGGACGUCUGG |
